# Modeling of enzyme-mediated glucose release to facilitate continuous feed in miniaturized cultivations

**DOI:** 10.1101/2023.05.14.540734

**Authors:** Annina Kemmer, Linda Cai, Stefan Born, M. Nicolas Cruz Bournazou, Peter Neubauer

## Abstract

When striving for maximal throughput at minimal volumes while cultivating close to industrial conditions, simple and robust feeding strategies offer important advantages. Enzyme-mediated glucose cleavage from dextrin is an easy way of imitating continuous fed-batch in the small scale, with no complex equipment required. While the release rate – and thus the feed rate – can be controlled by adapting the enzyme concentration, it strongly depends on the concentration of the involved substances and the environmental conditions. Thus, it is a challenge to use the technology for controlling the specific growth rate, as it is commonly done with feed pumps. For solving this problem, we present here a mathematical model that extends simple Michaelis-Menten kinetics by considering different substrate fractions and can be applied to control the glucose release rate even in high throughput experiments. The fitted model was used during automated microbial cultivations to control the growth rate in quasi-continuous fed-batch processes and to realize different exponential growth rates by intermittent additions of enzyme and dextrin by a liquid handling robot system. We thus present an approach for defined biocatalytically controlled glucose supply of small-scale systems, where – if at all – continuous feeding was only possible with low accuracy or high technical efforts until now.

## 1 Introduction

Fed-batch is the preferred cultivation mode for biotechnology processes to prevent substrate accumulation, by-product formation, or engineering limitations. As the performance of cells strongly depends on the cultivation mode, this technology should be already applied in the very early steps of process development, where often big numbers of parallel cultivations are performed e.g. for screening of preferable clones (Lara et al. 2006; Enfors et al. 2001).

Several approaches for fed-batch strategies in miniaturized systems have been developed in the recent years (Teworte et al. 2022). However, one difficulty still is the implementation of an accurate and simple continuous feeding technology. In miniaturized systems, if possible at all, the feeding of the carbon source, which in many cases is glucose, is often discontinuous – either by droplet formation at low pump rates or by bolus feeding through the pipetting needles of a liquid handling robot. These oscillations of substrate availability (Figure 1(b)) produce very relevant effects in the culture compared to continuous feeding (Figure 1(a)). Short periods of substrate excess followed by starvation have an influence on growth and product formation (Neubauer and Junne 2016; Delvigne et al. 2018). While controlled oscillations can be a valuable tool to mimic the concentration gradients present in industrial-scale bioreactors and their input on microbial physiology (Emmanuel Anane et al. 2019; E. Anane et al. 2019), they limit the comparability of small-scale and benchtop-scale cultivations.

**Figure 1.**
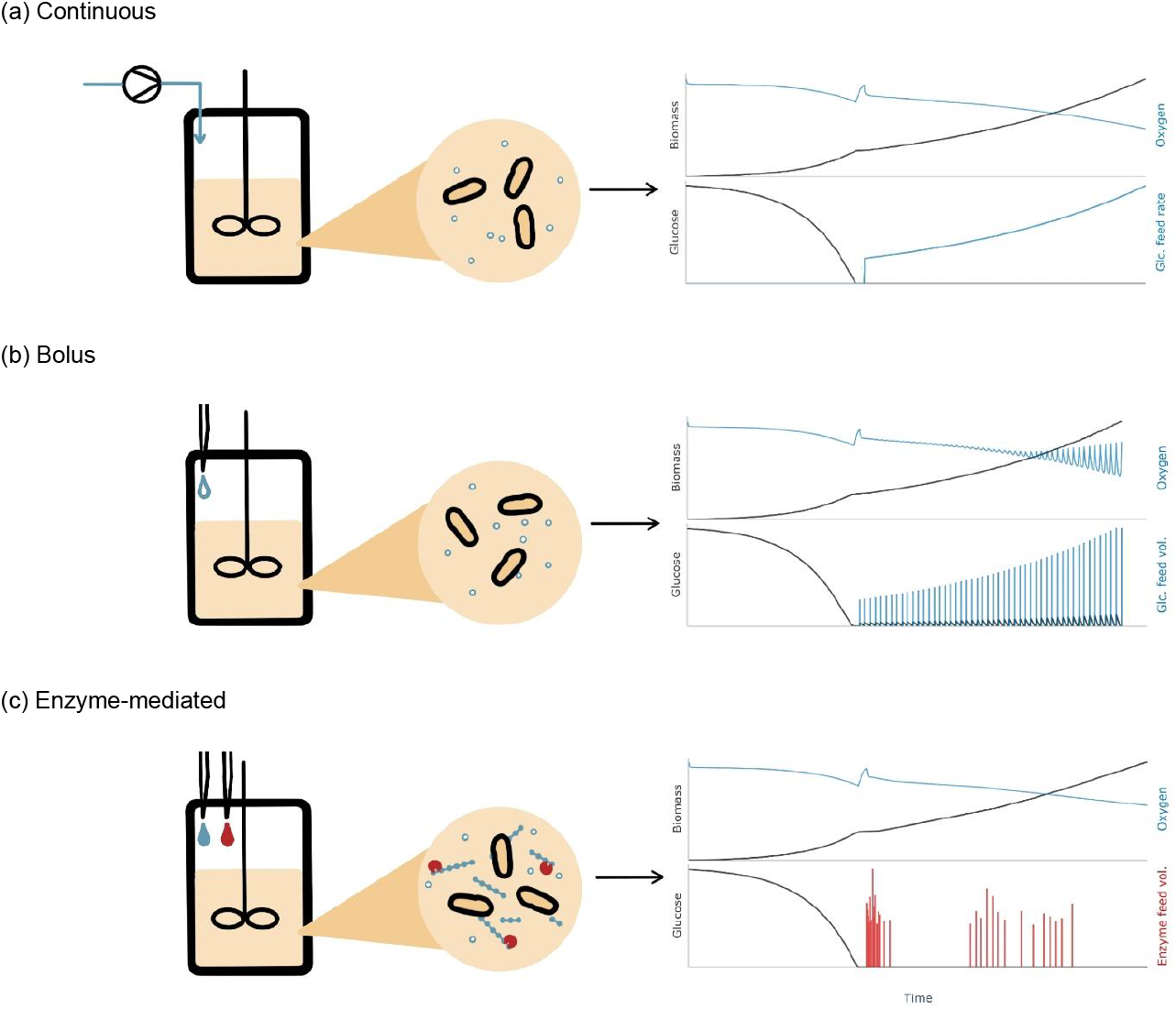
Comparison of (a) pumped, (b) bolus and (c) enzyme-mediated feed. In the larger scale, glucose feed is added continuously via pumps while in miniaturized systems discreet pulses are given. Continuous glucose feed in the small scale can be realized by enzyme-catalyzed glucose release from a dextrin. The glucose release rate is tuned by enzyme and dextrin additions by the needles of a liquid handling site.

Methods for continuous feed include miniaturized pumps (Gebhardt et al. 2011; Funke et al. 2010), glucose release by diffusion from a polymer matrix (Jeude et al. 2006; Scheidle et al. 2010; Habicher et al. 2020; Bähr et al. 2012) or enzyme-mediated glucose release from an insoluble (Panula-Perälä et al. 2008) or soluble polysaccharide (Krause et al. 2010). Miniaturized pumps offer the advantage of a direct scale-down from the benchtop techniques, while being hard to implement and high-maintenance. Regarding glucose release from a polymer matrix, the control of the feed rate is difficult, and its implementation in stirred systems is not straight forward, as gels or beads containing the substrate are added (Jeude et al. 2006; Scheidle et al. 2010). Enzyme-mediated glucose release (Krause et al. 2010; Panula-Perälä et al. 2008) describes the release of glucose monomers from starch or dextrins through hydrolysis of glycosidic bonds by glycolytic enzymes, such as glucoamylase.

Enzymatic glucose fed-batch has been successfully applied for expression of different difficult to express proteins in *Escherichia coli* in shake flasks and microwell plates such as single-domain antibodies (Zarschler et al. 2013), valinomycin synthetase (Li et al. 2014) or ribonuclease inhibitor (Šiurkus et al. 2010) as well as during online-adaptive experiments for growth characterization of *E. coli* in minibioreactors (MBRs) (Cruz Bournazou et al. 2017), and microtiter plate cultivations (Jansen et al. 2019).

The release rate, and thus the feed rate, depends on the concentrations of the reaction components. As all components are in a liquid state, this method offers the potential of being handled by liquid handlers. Continuous feeding of glucose with adjustable rates can therefore be achieved by bolus additions of enzyme and dextrin (Figure 1(c)).

However, the hydrolysis reaction represents a non-linear, highly complex system, which depends on various environmental conditions. Despite the concentrations of the enzyme, dextrin and glucose, the composition of the dextrin (Appl, Baganz, and Hass 2021), the pH (Hiromi, Takahashi, et al. 1966; Sauer et al. 2000) and the temperature (James and Lee 1997; Marc et al. 1983) potentially have an influence on the glucose release rate. Other impacting factors include the polysaccharide chain length (Ono, Hiromi, and Zinbo 1964), high viscosity of concentrated starch solutions (Miranda et al. 1991), branching (Sanromán, Murado, and Lema 1996), competition between different types of glycosidic bonds (Hiromi, Hamauzu, et al. 1966), different binding modes of the enzyme to the substrate (Hiromi, Ohnishi, and Tanaka 1983), and enzyme inactivation (Polakovič and Bryjak 2002).

Efforts to describe these dependencies with the help of mathematical models exist and could, if applied to the enzyme-mediated glucose release systems, control and predict the experiments. While for starch there are examples of data-driven models (Wojciechowski, Koziol, and Noworyta 2001; Bryjak et al. 2000), most commonly, mechanistic models with Michaelis-Menten type kinetics have been proposed (Ono, Hiromi, and Zinbo 1964; Miranda et al. 1991; Polakovič and Bryjak 2004; Sanromán, Murado, and Lema 1996). However, these models have been developed in the context of saccharification processes in the food industry, i.e., the production of high-glucose syrups under conditions which are not favorable for microbial cultivations. Thus, these models are not directly applicable.

In this study, we present a mechanistic model which has been selected to describe starch hydrolysis under the conditions relevant for microbial cultivations. The model is used to predict and plan the glucose release in miniaturized parallel cultivations. The release rate, and thus the feed rate, is adjusted by additions of enzyme and dextrin. We show that using this approach, substrate-limited exponential growth without oscillations in the substrate availability is achieved.

## 2 Material and Methods

### 2.1 General Experimental Set-up

All experiments were performed in the mL-scale in a high-throughput platform (Haby et al. 2019). The cultivation platform consists of a Tecan Freedom Evo 200 liquid handling site (LHS) (Tecan Group Ltd, Männedorf, Switzerland) hosting a bioREACTOR® 48 system (2mag AG, Munich, Germany) for up to 48 experiments.

If not stated differently, the experiments, with and without cells, were conducted in mineral salt medium (MSM) with varying initial glucose and dextrin concentrations. For the MSM medium, the following solutions were prepared and sterilized separately: basic salt solution (autoclaved, prepared as 5x concentrate) (final medium concentration per L): 12.6 g K_2_HPO_4_, 3.6 g NaH_2_PO_4_ × 1H_2_O, 2 g Na_2_SO_4_, 2.468 g (NH_4_)_2_SO_4_, 0.5 g NH_4_Cl, 1 g (NH_4_)_2_-H-citrat; dextrin concentrate (autoclaved) (per L): 150 g maltodextrin; magnesium sulfate concentrate (autoclaved) (per L): 246.48 g MgSO_4_ × 7H_2_O; thiamine concentrate (sterile filtered) (per L): 50 g thiamine hydrochloride; trace element solution (sterile filtered) (5x) (per L): 0.5 g CaCl_2_ × 2H_2_O, 0.18 g ZnSO_4_ × 7H_2_O, 0.1 g MnSO_4_ × H_2_O, 20.1 g Na_2_-EDTA, 16.7 g FeCl_3_ × 6H_2_O, 0.16 g CuSO_4_ × 5H_2_O, 0.725 g Ni(NO_3_)_2_ × 6H_2_O; and D-glucose concentrate (autoclaved) (per L): 600 g D-glucose (C_6_H_12_O_6_). For 1 L final medium 200 mL basic salt solution, 2 mL magnesium sulfate concentrate, 2 mL thiamine concentrate, 2 mL trace element solution, and appropriate volume of dextrin and glucose concentrate, and deionized water were combined to reach desired dextrin and glucose concentrations and the target end volume. The pH was adjusted using 7 M ammonia. The glucose release was achieved by addition of the enzyme glucoamylase stock solution (3000 U L^-1^).

Sample plates were prepared by adding 15 μL of 2 M NaOH_(aq)_ to each well of 96-well V-bottom deep well plates (Corning Inc., New York, USA). The plates were dried at 80 °C for at least 24 h. Samples of 200 μL were taken automatically by the pipetting tips of the Tecan LHS, pipetted into chilled sample plates and mixed to dissolve the dried sodium hydroxide. This results in an increase of the pH of up to > pH 13 which stops the glucose release by inactivating the enzyme, resp. inhibits cell activity (Nickel et al. 2016). Immediately after, the stopped samples of the cell-free release experiments and the microbial fed-batch cultivations were analyzed as described below.

Setpoints for enzyme, glucose and dextrin additions, pH and temperature control, as well as all measurements were stored in a central database, which allows rapid data analysis.

### 2.2 Cell-free Experiments for Selection of the Glucose Release Model

Under sterile conditions, each MBR was filled with 11 mL of the MSM at pH 7.0 containing different initial concentrations of dextrin and glucose (Table 1). The kinetic experiments were started by enzyme addition to each MBR by the pipetting tips of the Tecan LHS. After approx. 6 h, dextrin and glucose pulses were added to some experiments to evaluate the possibility of substrate and product inhibition. A detailed overview of the experimental conditions is given in the supplementary material (Table S1). The cultivations were aerated with 0.4 L_n_ min^-1^ (mass flow at 0 °C, 1.013 bar) pressurized air, the temperature was set to 30 °C, and agitation was set to 1600 rpm. Evaporation at the cultivation conditions was determined to be approx. 30 L h^-1^.

**Table 1.** Design variables of the enzymatic release experiments. The initial concentration of dextrin *D*_0_, glucose *G*_0_ and enzyme *E*_0_ were varied. To some experiments, dextrin *D*_*add*_ and glucose *G*_*add*_ was added after approx. 6h.

| Parameter | Description | Unit | Values |
| --- | --- | --- | --- |
| $D_0$ | Initial dextrin concentration | g L<sup>-1</sup> | 15, 30 |
| $G_0$ | Initial glucose concentration | g L<sup>-1</sup> | 0, 3.75, 7.5, 15 |
| $E_0$ | Initial enzyme concentration | U L<sup>-1</sup> | 10, 20 |
| $D_{add}$ | Dextrin additions | g L<sup>-1</sup> | 0, 5.25, 10.5 |

Samples were taken every 1 to 2 h before and every 2 to 4 h after addition of glucose or dextrin into the sample plates with dried stop solution. The glucose concentration was measured directly in the sample using a Cedex Bio HT Analyzer (Roche Holding AG, Basel, Switzerland).

### 2.3 Enzymatic Fed-batch Experiments

MBR cultivations were performed with *Escherichia coli* BL21(DE3), carrying the plasmid pET28-NMBL-eGFP-TEVrec-(*V*_2_*Y*)_15_-His, expressing a recombinant fusion protein of ELP and eGFP, under the isopropyl-β-D-thiogalactopyranosid (IPTG) inducible *lac*UV5-promoter. Detailed information about the plasmid can be found in Huber et al. (2014) and Schreiber et al. (2019). For a first preculture, 10 mL LB medium, containing 16 g L^-1^ tryptone, 10 g L^-1^ yeast extract, 5 g L^-1^ NaCl, was supplemented with 10 μL Kanamycin stock (50 mg mL^-1^) and inoculated with 1 mL cryostock. The preculture was cultivated in a 125 mL UltraYield® Flask (Thomson Instrument Company, Oceanside, USA) sealed with an AirOtop® enhanced flask seal (Thomson Instrument Company, Oceanside, USA) for 5 h at 37 °C and 220 rpm in an orbital shaker (50 mm amplitude) (LT-X, Adolf Kühner AG, Birsfelden, Switzerland) until an OD_600_ approx. 7. For the second preculture, 25 mL EnPresso® B (EnPresso GmbH, Berlin, Germany) with 2.5 μL antifoam and 25 μL kanamycin stock (50 mg mL^-1^) in a 250 mL UltraYield flask were inoculated with the first preculture to an OD_600_ of 0.25. Glucose release was started by addition of 12.5 μL glucoamylase to obtain 6 U L^-1^. The flask was sealed with an AirOTop enhanced flask seal and cultivated at 37 °C and 220 rpm for 14 h.

Each MBR was filled with MSM (see section *2.1* *General Experimental Set-up*) and the second preculture was added automatically by the liquid handling needles of the Tecan LHS to an OD_600_ of 0.25, corresponding to a biomass concentration of ∼0.0925 g L^-1^, in a total of 10 mL medium. An overview of the experimental conditions can be found in Table 2. 24 MBR experiments were performed, including enzymatic feed with a low and a slightly higher exponential feed as well as experiments with only two enzyme additions but a similar target release. Two types of controls were included: 1) Cultivations with bolus feed with the same feed rate as during the enzymatic feed experiments, and 2) cell-free release experiments to verify that the previously fitted glucose release model is valid.

**Table 2.** Experimental conditions of the enzymatic fed-batch experiment. Each condition was performed in triplicates. The conditions differed in their feed methods and set growth rates (before and after induction).

| Condition | Feed method | $\mu_{set}$ [h<sup>-1</sup>] | $G_0$ [g L<sup>-1</sup>] | $D_0$ [g L<sup>-1</sup>] | Additions | Cells |
| --- | --- | --- | --- | --- | --- | --- |
|  |  | Before ind \| after ind |  |  |  |  |
| 1 – control pulse | Bolus feed | 0.18 \| 0.09 | 5 | 2 | Glucose | Yes |
| 2 – control pulse | Bolus feed | 0.22 \| 0.11 | 5 | 2 | Glucose | Yes |
| 3 – enzymatic feed | Enzymatic release | 0.18 \| 0.09 | 5 | 40 | Dextrin, Enzyme | Yes |
| 4 – enzymatic feed | Enzymatic release | 0.22 \| 0.11 | 5 | 40 | Dextrin, Enzyme | Yes |
| 5 – enzymatic feed | Enzymatic release (two additions) | 0.18 \| 0.09 | 5 | 40 | Dextrin, Enzyme | Yes |
| 6 – enzymatic feed | Enzymatic release (two additions) | 0.22 \| 0.11 | 5 | 40 | Dextrin, Enzyme | Yes |
| 7 – control cell-free | Enzymatic release | 0.18 \| 0.09 | 0 | 40 | Dextrin, Enzyme | No |
| 8 – control cell-free | Enzymatic release | 0.18 \| 0.09 | 0 | 40 | Dextrin, Enzyme | No |

The cultivations were aerated with 5 L_n_ min^-1^ pressurized air, the temperature was set to 30°C, and agitation was set to 2800 rpm. The cultivations were controlled at the target pH of 7.0 +/- 0.2 by addition of 7 M ammonia or 3 M phosphate solution, respectively. Evaporation at the cultivation conditions was determined to be approx. 30 L h^-1^. After the batch phase of approx. 9 h, the “control pulse” cultivations were supplied with bolus additions of a 377.3 mg L^-1^ glucose feed by the robotic pipetting tips (Kemmer et al. 2022). For the remaining MBRs, the glucose release, and thus the glucose feed, was initiated and controlled by repeated enzyme additions by the liquid handler. Additionally, 40 μL of the dextrin stock solution were added every 10 min to keep the dextrin concentration high. Production of the recombinant protein was induced 6 h after fed-batch start by addition of 0.5 M IPTG stock to a concentration of 0.2 mM.

DOT and pH were measured online every 30 to 60 sec by fluorescence sensors (PreSens Precision Sensing GmbH, Regensburg, Germany). At-line samples were taken every 1 to 2 h starting approximately 1 h before batch end. The optical density at 600 nm (OD_600_) was measured in a Synergy MX microwell plate reader (BioTek Instruments GmbH, Bad Friedrichshall, Germany) after manual sample dilution with 0.9 % saline solution. The cells were separated from the supernatant by centrifugation at 15000*g* for 10 min, and the glucose, acetate and magnesium concentrations were determined in the supernatant using the Cedex Bio HT Analyzer. An additional estimate of the biomass concentration 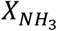 was obtained from the added ammonia as described by Kemmer et al. (2022).

### 2.4 Dynamic Modeling

Mechanistic modeling was performed using an in-house developed Python-based framework for simulation using symbolic differentiation (Andersson, Åkesson, and Diehl 2012), and parameter estimation using lmfit (Newville et al. 2014) and pygmo (Biscani and Izzo 2020). For lmfit the method “least_squares”, and for pygmo the algorithm “differential evolution” (“de1220”) with a population size of 20 and generation number of 200 was used. Samplings and inputs, namely the addition of dextrin, glucose, enzyme, and pH control reagents acid and base, are given as boluses and therefore represent discontinuous actions. After each of these actions, the simulation is stopped, the volume-dependent states are recalculated, and the simulation is restarted. The values of the model parameters were estimated as described by Kemmer et al. (2022).

## 3 Results

### 3.1 Selection and Parameter Fitting of the Glucose Release Model

The enzyme-mediated glucose release is based on the cleavage of D-glucose molecules from dextrin by the exo-acting enzyme glucoamylase (1,4-α-D-glucan glucohydrolase, EC 3.2.1.3) from *Aspergillus niger*. The enzyme catalyzes the hydrolysis of α-1-4 and α-1-6 glycosidic bonds from the non-reducing ends of starch and related poly- and oligosaccharides (Hiromi, Hamauzu, et al. 1966).

The kinetics of dextrin hydrolysis to glucose molecules by glucoamylase were investigated under the influence of various levels of the design variables glucose, dextrin, and enzyme concentration. To access the possibility of substrate and product inhibition, the influence of dextrin and glucose additions 6 h after initiation of glucose release was evaluated. The release reaction is dependent on the pH (Hiromi, Takahashi, et al. 1966; Sauer et al. 2000) and the temperature (James and Lee 1997). Thus, the experiments were controlled at 30 °C and pH 7.0, which are favorable conditions during microbial cultivations.

Initial model fitting approaches confirmed previous suppositions (Polakovič and Bryjak 2004; Hiromi, Hamauzu, et al. 1966; Sanromán, Murado, and Lema 1996) that the glucose release cannot be described using simple Michaelis-Menten type kinetics, or modifications of these kinetics for product inhibition or enzyme deactivation. Our data at pH 7 and 30 °C suggests that a part of the substrate was not susceptible to the enzymatic hydrolysis or at a much slower rate. This was in agreement with the findings of Polakovič and Bryjak (Polakovič and Bryjak 2004) who distinguish two fractions of starch, with one fraction much less susceptible to enzymatic hydrolysis.

Data of 24 release experiments was used simultaneously to fit the parameters of the two substrates models (Figure 2). However, including inhibition terms did not improve the model fit and was thus omitted. The model thus consists of the differential algebraic equations of the substrate dextrin *S* [g L^-1^], product glucose *P* [g L^-1^], the relative amount of susceptible substrate *W*_*S*_ [g g^-1^], enzyme *E* [U L^-1^] and the volume *V* [L]. Due to the conversion of anhydroglucose in starch into free glucose (J. P. Wang et al. 2006), a conversion factor of 1.111 is incorporated in equation 1:

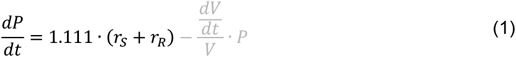

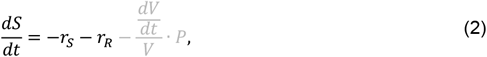

with the susceptible (S) substrate fraction *S*_*S*_ = *S*·*W*_*S*_ and the resistant (R) fraction *S*_*R*_ = *S* − *S*_*S*_.

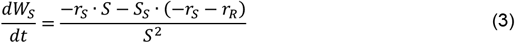

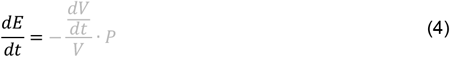

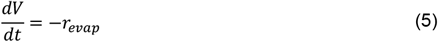

**Figure 2.**
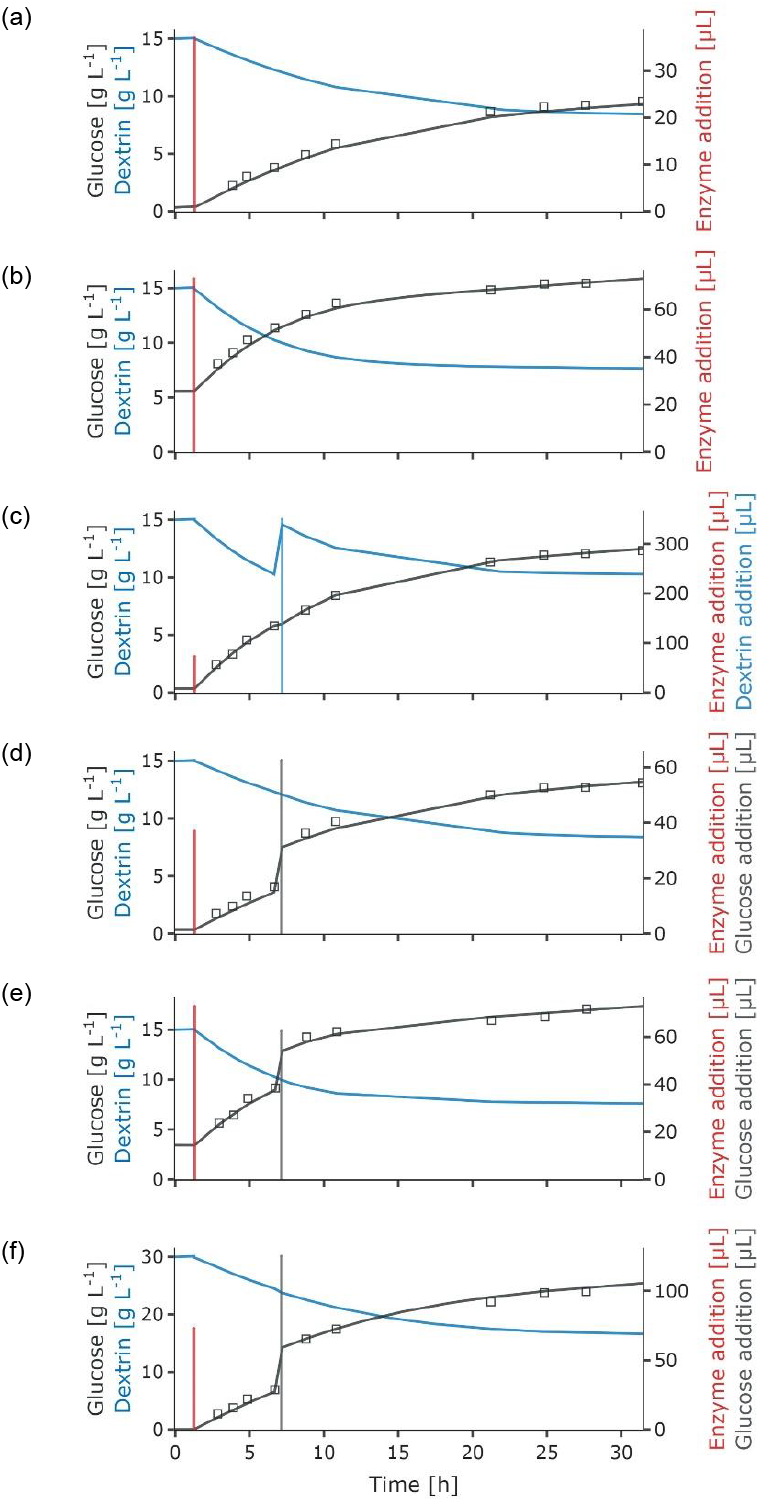
Glucose release experiments under different conditions regarding enzyme (E), glucose (G), and dextrin (D) concentration, specifically in (a) 15 gL^-1^ *D*_0_ and 10 UL^-1^ *E*_0_, (b) 15 gL^-1^ *D*_0_, 5.6 gL^-1^ *G*_0_ and 20 UL^-1^ *E*_0_, (c) 15 gL^-1^ *D*_0_, 10 UL^-1^ *E*_0_ and 5.25 gL^-1^ *D*_*add*_, (d) 15 gL^-1^ *D*_0_, 10 UL^-1^ *E*_0_ and 3.75 gL^-1^ *G*_*add*_, (e) 15 gL^-1^ *D*_0_, 3.5 gL^-1^ *G*_0_, 20 UL^-1^ *E*_0_ and 5.25 gL^-1^ *G*_*add*_, and (f) 30 gL^-1^ *D*_0_, 20 UL^-1^ *E*_0_ and 7.5 gL^-1^ *G*_*add*_. The glucose (measurements – black dots, simulation – black line) and dextrin concentration (simulation – blue line) are shown, as well as the added enzyme (red vertical line), glucose (grey vertical line), and dextrin volumes (light blue vertical line).

The release rates *r*_*S*_ [g (Lh)^-1^] for susceptible substrate and *r*_*R*_ for resistant substrate are calculated from the enzyme concentration *E* [g L^-1^], the catalytical constants *k*_*s*_ [g (UL)^-1^] and *k*_*R*_, and the Michaelis constant *K* [g L^-1^]:

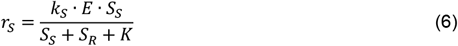

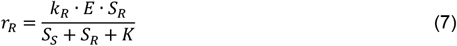

During incubation without dextrin the glucoamylase was found to be stable (data not shown). Thus, not degradation of the enzyme stock solution was assumed.

The determined parameters are shown in Table 3. Due to the comparatively high dextrin concentrations in the experiments the Michaelis constant *K* could not be fitted and was set to the value reported by Polakovič and Bryjak (Polakovič and Bryjak 2004). After 30 h reaction time a high amount of dextrin remains present, corresponding to the resistant fraction. This is in contrast to preliminary experiments where the polymer distribution during the release was characterized at pH 4 using gel permeation chromatography (GPC) (data not shown). Here, the reaction proceeded much faster and the difference between the two fractions of substrate was less pronounced.

**Table 3.** Fitted parameters of the enzymatic glucose release model with standard deviation.

| Parameter | Unit | Value | Standard deviation |
| --- | --- | --- | --- |
| $K$ | $\text{g L}^{-1}$ | 0.406 | (fixed) |
| $k_S$ | $\text{g (U h)}^{-1}$ | 0.13393567 | 0.00434840 (3.25 %) |
| $k_R$ | $\text{g (U h)}^{-1}$ | 0.00211669 | $4.5889 \cdot 10^{-4}$ (21.68 %) |
| $W_{S,0}$ | $\text{g g}^{-1}$ | 0.46419034 | 0.01071653 (2.31 %) |

### 3.2 Adaptation of the enzymatic glucose feed

For the enzymatic glucose feed experiments, the enzyme feed is determined by iterative comparison of the target glucose feed rate and the current glucose release rate. Enzyme additions were planned and executed by the liquid handler to control the glucose release rate and thus the set growth at the setpoint (see Table 2). The target glucose release rate is calculated as the difference between the current glucose concentration *P*, the target glucose concentration at a future timepoint *P*_*target*_ and the time difference from the current time to this future timepoint Δ*t*_*target*_. The enzymatic feed was calculated under the simplifying assumption that for the feeding intervals, difference quotients approximate the derivatives sufficiently well:

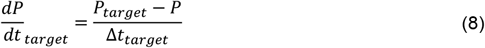

The necessary enzyme feed to achieve 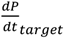 can be calculated from equation 2:

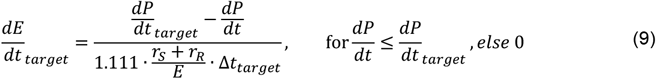

### 3.3 Application of Enzyme-mediated Glucose Release in Microbial Fed-Batch Cultivations

The results from the kinetic experiments were validated in parallel high-throughput mini bioreactor cultivations with enzymatic glucose feed. 24 experiments were performed, including control experiments with pulsed feed as well as cell-free release experiments. Glucose release was initiated by enzyme addition after the batch phase of approx. 10 h. Throughout the fed-batch phase, more enzyme was added to control the glucose release at the desired setpoint. Additionally, dextrin was added to maintain a high substrate concentration. As in the cell-free release experiments, the cultivations were performed at 30 °C and pH 7. However, higher volumes of enzyme (factor 2-3) were added.

A combined mechanistic model including (i) the previously described enzyme-mediated glucose release and (ii) the microbial growth (Kemmer et al. 2022) was fitted to the validation experiments. First, the parameters of the cell-free experiments were reassessed. The glucose release was slightly overestimated when using the parameters obtained in the model calibration experiments. As shown in Figure 3a, a good fit of the glucose can be achieved with a lower catalytical constant for the susceptible substrate *k*_*s*_ = 0.076 (*UL*)^−1^. The simulated concentrations of the susceptible and resistant dextrin fractions show an accumulation of the resistant dextrin with increased experimental time. This is due to the extremely low hydrolysis of the resistant substrate and simultaneous addition of dextrin stock during the feeding phase to maintain a sufficient concentration of susceptible substrate.

**Figure 3.**
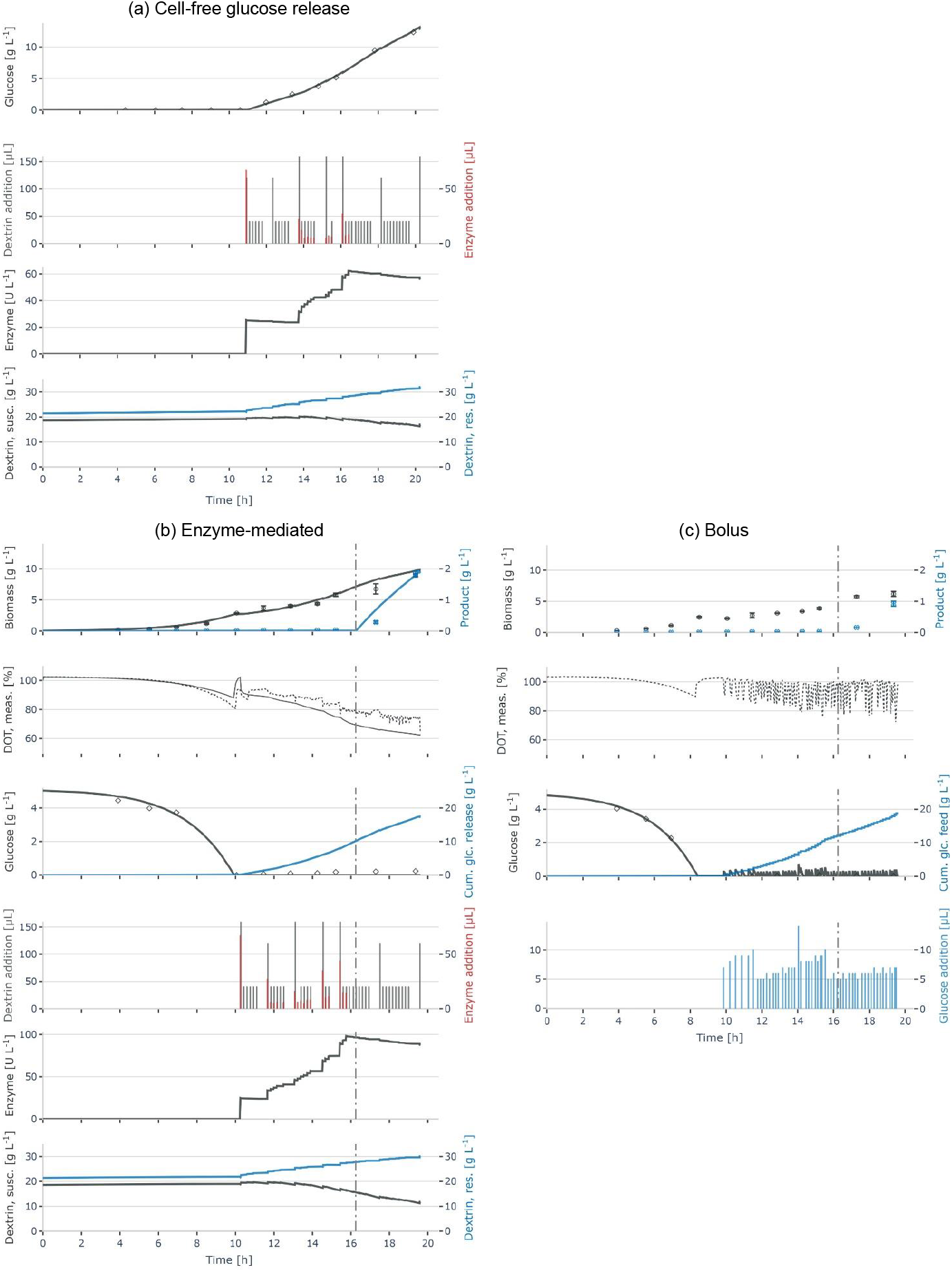
Application of mechanistic glucose release model to facilitate continuous feeding in *E. coli* cultivations. Glucose is released continuously by enzyme-mediated hydrolysis of dextrin and (a) accumulates in cell-free control experiments, or (b) is metabolized by the cells of a microbial cultivation. In (c) cell growth during a cultivation with bolus additions of glucose feed is shown as comparison. Model simulations are shown as lines, measurements as symbols or dotted line (DOT), and inputs as bars. Besides the glucose measurements and simulations, the glucose which is either (b) released or (c) added during the feeding is shown.

In a next step, with fixed glucose release parameters, the growth-related model parameters were estimated using the data of the experiments with enzyme-mediated glucose feed (see Table S2 in the supplementary material). The model can describe the course of the biomass, product and glucose data with sufficient accuracy (Figure 3b). The oxygen (DOT) profile shows a continuous decrease, indicating that a continuous substrate release, and thus consumption, with increasing rate is achieved.

Conversely, in the bolus-fed control (Figure 3c) oscillations in the oxygen signal can be observed as cells switch between excess and starvation in between the glucose additions. While the amount of glucose which is added is similar to the released glucose in the enzyme-mediated glucose fed cultivations, the achieved amount of biomass at the end of the fed-batch phase is significantly smaller. As the influence cannot be quantified yet, the simulations for biomass, product and oxygen are not shown here.

## 4 Discussion

Data from cell-free glucose release experiments was successfully fitted using a Michaelis-Menten kinetic extended by the assumption of two different substrates, a fraction which is more susceptible to hydrolysis and a resistant one (Polakovič and Bryjak 2004). Glucose cleavage from the two assumed to types of dextrin was catalyzed by the amylolytic enzyme glucoamylase.

The catalytical constants for release of glucose from the susceptible substrate *k*_*S*_ and from the resistant substrate *k*_*R*_ were found to be smaller than stated in the original publication (Polakovič and Bryjak 2004), indicating a lower glucose release. Additionally, the initial relative amount of susceptible substrate *W*_*S*,0_ was found to be 46 %, which is lower than the 77 % stated in previous publications (Polakovič and Bryjak 2004; Zanin and Moraes 1996). This is reasonable as the hydrolysis reaction is pH-dependent (Sauer et al. 2000) and, with a pH of 7.0 and a temperature of 30° C, the experimental conditions in this study are far from the optimal conditions of the enzyme glucoamylase (Dalmia and Nikolov 1991). Further analysis of the two fractions would be necessary for a quantitative explanation of the behavior. Preliminary analysis of the polymer distribution during the release at pH 4 show a faster reaction rate and less differences between the two substrate fractions (data not shown here). This supports the assumption that this change in behavior is at least partly due to the different pH values. Our data did not allow an accurate estimation of the catalytical constants for the less susceptible fraction *k*_*R*_, which would need much longer reaction times than we could achieve, and also is not relevant for the considered fed-batch cultivations. Product inhibition (Hiromi, Ohnishi, and Tanaka 1983; Polakovič and Bryjak 2004) and substrate inhibition (Miranda et al. 1991; Sanromán, Murado, and Lema 1996) do not seem to play a role in our application.

The aim of this study was the selection of a model under conditions which are suitable for microbial cultivations, thus a temperature of 30° C and pH of 7 were chosen. The fitted model describes the data of the glucose release as well as the microbial growth well. Microbial cultivations are executed at varying temperatures of ∼25 – 37 °C and pH values of 6.5 – 7.1 for *Escherichia coli* or 5.8 – 6.5 for yeast. To facilitate the use of the glucose release model in a broader application spectrum, the influence of the temperature and the pH should be studied and additionally included in the model. For a constant temperature and pH value, the model parameters can be adapted based on experimental data and used for prediction of the glucose release without the need to change the model structure. However, for experimental settings including dynamic changes in the temperature and pH, the influence of these factors on the release rate must be included in the model equations, as it has been done for e.g., α-amylase (Hiromi, Takasaki, and Ono 1963) as well as ligninolytic and cellulolytic enzymes (G. Wang et al. 2012).

The glucose release model was combined with a macro-kinetic model describing the growth of *Escherichia coli* in high-throughput enzyme-mediated glucose feed experiments. Data of cell-free glucose release control experiments showed that *k*_*s*_ estimated to be 0.0756 g (U h)^-1^ and thus smaller than the 0.134 g (U h)^-1^ estimated in the previous model identification experiments. Consequently, the attained glucose release rates were lower than the setpoints (Table 2). The reasons for this might be differences between substrate batches, resulting a different substrate composition, e.g., in terms of the length. A higher agitation and aeration can lead to inactivation of the enzyme and thus a lower activity (Maa and Hsu 1997). Additionally, a negative effect on the reaction rate has been observed due to limitations in the mass transfer at higher substrate concentrations (Sanromán, Murado, and Lema 1996), which is the case in the enzymatic fed-batch experiments.

The main aim to apply enzyme-mediated glucose release in this study was avoiding oscillations in the glucose availability. The oxygen signal of the cultivations fed by glucose release shows a mainly smooth, steadily decreasing course. This confirms that continuous substrate release with an increasing release rate has been achieved. Conversely, the oxygen signal of the bolus-fed cultivations shows the peaks which are characteristic for the repeated switch between maximum growth after the pulse has been given, and starvation upon consumption of the glucose shortly after.

In cultivations with enzyme-mediated feed, the simulated oxygen signal deviates from the measured one, with the oxygen consumption being underestimated in batch phase and overestimated in fed-batch phase. While the cause for this phenomenon cannot be deduced from the results, it might lie in the influence of dextrin on viscosity (Miranda et al. 1991), mass transfer (Sanromán, Murado, and Lema 1996), and thus on oxygen transfer, which changes with the depolymerization of the substrate during the reaction (Bryjak et al. 2000).

The bolus-fed cultivations show a significantly lower biomass growth than the continuously fed cultures despite comparable glucose supply. This is the typical well-described response of many microbial cells to the oscillating feast vs. famine availability of glucose (feed zone effect) (Neubauer and Junne 2010). Our results thus suggest that the influence of oscillations can be studied in these small-scale systems. This confirms the potential use of miniaturized cultivations to study scale effects (Anane et al. 2019), and would be valuable to include in further studies.

Our results show that continuous glucose feeding with defined release rates is possible in small-scale systems. However, accumulation of resistant dextrin and low release rates set limits in the possible length and setpoints for the release rate. Here, the addition of debranching enzymes, cleaving the α-1-6 glycosidic bonds, (Guzmán-Maldonado, Paredes-López, and Biliaderis 2009) could improve the release rate. This, however, would require the incorporation of the respective kinetics into the mechanistic model.

Enzymatic glucose feed has been previously applied in microscale experiments to regulate the growth rate (Jansen et al. 2019). The employed method presents a straightforward control approach but does not allow to distinguish between different factors that impact growth, namely lack of enzyme, lack of susceptible substrate or cellular factors. A combination with our model-based approach would enable us to do this. In order to cope with model uncertainties and measurement errors, the development of a state estimator (e.g., a variant of the Kalman filter) is desirable. This would increase the reliability of the model predictions and of the planned feeds.

## 5 Conclusions

We present an approach for defined continuous glucose feeding in small scale systems. A glucose release model considering two substrate fractions was successfully fitted to experimental data under different conditions relevant to microbial cultivations. Subsequently, we demonstrated the usage of enzymatic glucose feed in validation experiments with *Escherichia coli*. Different exponential growth rates could be achieved by intermittent additions of enzyme and dextrin during enzyme-mediated glucose feed experiments. This approach mimics industrial feeding strategies and thus enables small-scale screening of potential production strains and cultivation settings close to industrial conditions. To facilitate the use of the glucose release model in a broader application spectrum, the influence of the temperature and the pH must be included in the model.

## Author Contributions

AK, NC, SB and PN contributed to the conception and design of the study. AK and LC performed the experiments and analytics. AK wrote the software and performed the data curation. AK and SB performed the fitting of model parameters. AK and LC prepared the figures. PN and NC provided resources and funding. AK wrote the first draft of the manuscript. All authors contributed to manuscript revision.

## Acknowledgments

We thank Prof. Dr. Stefan Schiller and Dr. Matthias Huber from Johann Wolfgang Goethe-Universität Frankfurt am Main for providing the *Escherichia coli* strain. We also thank Dr. Jens Buller and Dr. Hendrik Wetzel from Fraunhofer IAP for the GPC analysis.

## Data Availability

The original contributions presented in the study are publicly available. This data can be found here: https://git.tu-berlin.de/bvt-htbd/public/kemmer_2022_enzymatic-feed.git.

