## Supplementary material for "Modeling of enzyme-mediated glucose release to facilitate continuous feed in miniaturized cultivations"

### Supplementary Data

### Experimental conditions for model calibration experiments

A detailed overview of the experimental conditions of the cell-free glucose release experiments is given in Table S1.

**Table S1.** Detailed overview on the experimental conditions of the 24 enzymatic release experiments. The initial concentration of dextrin $D_0$, glucose $G_0$ and enzyme $E_0$ were varied. To some experiments, dextrin $D_{add}$ and glucose $G_{add}$ was added after ~ 6h.

| Exp. | $D_0$ [g L$^{-1}$] | $G_0$ [g L$^{-1}$] | $E_0$ [U L$^{-1}$] | $D_{add}$ [g L$^{-1}$] | $G_{add}$ [g L$^{-1}$] |
|---|---|---|---|---|---|
| 1 | 15 | 0 | 10 | - | - |
| 2 | 15 | 0 | 20 | - | - |
| 3 | 15 | 0 | 10 | - | 3.75 |
| 4 | 15 | 0 | 20 | - | 3.75 |
| 5 | 15 | 0 | 10 | 5.25 | |
| 6 | 15 | 0 | 20 | 5.25 | |
| 7 | 30 | 0 | 10 | - | - |
| 8 | 30 | 0 | 20 | - | - |
| 9 | 15 | 7.5 | 10 | - | - |
| 10 | 15 | 7.5 | 20 | - | - |
| 11 | 30 | 15 | 10 | - | - |
| 12 | 30 | 15 | 20 | - | - |
| 13 | 30 | 0 | 10 | - | 7.5 |
| 14 | 30 | 0 | 20 | - | 7.5 |
| 15 | 30 | 0 | 10 | 10.5 | - |
| 16 | 30 | 0 | 20 | 10.5 | - |
| 17 | 15 | 3.75 | 10 | - | - |
| 18 | 15 | 3.75 | 20 | - | - |
| 19 | 15 | 3.75 | 10 | - | 3.75 |
| 20 | 15 | 3.75 | 20 | - | 3.75 |
| 21 | 30 | 7.5 | 10 | - | - |
| 22 | 30 | 7.5 | 20 | - | - |
| 23 | 30 | 7.5 | 10 | - | 7.5 |
| 24 | 30 | 7.5 | 20 | - | 7.5 |

### Enzymatic fed-batch experiments

Table S2 contains the constants which were used, and parameters which were obtained from parameter fitting of the enzymatic fed-batch experiments. The full code is given in: https://git.tu-berlin.de/bvt-htbd/public/kemmer_2022_enzymatic-feed.git.

**Table S2.** Overview on parameters and constants. Description of parameters and constants which were obtained in the model fitting enzymatic fed-batch experiments are given, including the origin of the value.

| Parameter | Description | Unit | Value | Origin / source |
|---|---|---|---|---|
| ***E. coli* growth model:** | | | | |
| $q_{s,max}$ | Maximum specific substrate uptake rate | $gg^{-1}h^{-1}$ | 1.342 | Data fitting |
| $q_m$ | Specific maintenance coefficient | $gg^{-1}h^{-1}$ | 0.0519 | Data fitting |
| $q_{Ac,max}$ | Maximum specific acetate consumption rate | $gg^{-1}h^{-1}$ | 1.554 | Data fitting |
| $q_{Ap,max}$ | Maximum specific acetate production rate | $gg^{-1}h^{-1}$ | 0.186 | Data fitting |
| $K_S$ | Affinity constant for substrate (glucose) consumption | $gL^{-1}$ | 0.001 | Data fitting |
| $K_{qS}$ | Monod-type saturation constant for intracellular acetate production, dependent on the intracellular substrate flux $q_S$ | $gg^{-1}h^{-1}$ | 3.872 | Data fitting |
| $K_A$ | Affinity for acetate consumption | $gL^{-1}$ | 0.420 | Data fitting |
| $K_{i,AS}$ | Inhibition of acetate uptake by glucose | $gL^{-1}$ | 0.929 | Data fitting |
| $Y_{AS,of}$ | Yield of acetate on substrate (overflow metabolism) | $gg^{-1}$ | 0.667 | Data fitting |
| $Y_{XS,em}$ | Yield of biomass on substrate, exclusive maintenance | $gg^{-1}$ | 0.518 | Data fitting |
| $Y_{XA}$ | Yield of biomass on acetate, exclusive maintenance | $gg^{-1}$ | 0.356 | Data fitting |
| $Y_{\mathrm{OS}}$ | Oxygen used per gram of glucose metabolized | $gg^{-1}$ | 1.067 | Stoichiometric constant (Xu, Jahic, and Enfors 1999) |
| $Y_{\mathrm{OA}}$ | Oxygen used per gram of acetate metabolized | $gg^{-1}$ | 1.067 | Stoichiometric constant (Xu, Jahic, and Enfors 1999) |
| $Y_{\mathrm{PS}}$ | Yield of product on substrate | $gg^{-1}$ | 2.00 | Data fitting |
| $d_{Sox,P}$ | Distribution constant defining fraction of $q_{Sox}$ going into product formation | $\%100^{-1}$ | 0.123 | Data fitting |
| **Glucose release model:** | | | | |
| $K$ | Michaelis constant | $g\ L^{-1}$ | 0.406 | (Polakovič and Bryjak 2004) |
| $k_S$ | Catalytical constant for the susceptible substrate | $g\ (U\ h)^{-1}$ | 0.0756 | Data fitting |
| $k_R$ | Catalytical constant for the resistant substrate | $g\ (U\ h)^{-1}$ | 0.00212 | Data fitting |
| $W_{S,0}$ | Initial relative amount of susceptible substrate | $g\ g^{-1}$ | 0.464 | Data fitting |

| Constant | Description | Unit | Value | Origin / source |
|---|---|---|---|---|
| $H$ | Henry constant | % L $g^{-1}$ | 14000 | Physical constant |
| $\tau$ | Response time of the oxygen sensor | h | 65 | Experimental data |
| $k_L a$ | Volumetric oxygen transfer coefficient | $h^{-1}$ | 280 | Data fitting |
| $DOT^*$ | Dissolved oxygen concentration at saturation (aeration with 5 $Lmin^{-1}$ air) | % | 103 | Experimental data |
| $T$ | Temperature | ° C | 30 | Experimental data |
| $r_{evap}$ | Evaporation rate | $Lh^{-1}$ | 0.0005 | Experimental data |
| $S_{i,glc}$ | Glucose concentration in the feed | $gL^{-1}$ | 377.3 | Input concentration |
| $S_{i,enz}$ | Enzyme concentration in the feed | $UL^{-1}$ | 3000 | Input concentration |
| $S_{i,dex}$ | Dextrin concentration in the feed | $gL^{-1}$ | 115 | Input concentration |
| $C_A$ | Carbon concentration in acetate | mol C $g^{-1}$ | 0.033 | Chemical formula (Xu, Jahic, and Enfors 1999) |
| $C_S$ | Carbon concentration in the substrate (glucose) | mol C $g^{-1}$ | 0.033 | Chemical formula (Xu, Jahic, and Enfors 1999) |
| $C_X$ | Carbon concentration in the biomass | mol C $g^{-1}$ | 0.041 | Stoichiometric analysis (Xu, Jahic, and Enfors 1999) |
| $C_P$ | Carbon concentration in the product | mol C $g^{-1}$ | 0.044 | Stoichiometric analysis |
